# Persistence of Zika virus RNA in the epididymis of the male reproductive tract

**DOI:** 10.1101/2020.10.08.330019

**Authors:** Megan B. Vogt, Francesca Frere, Seth A. Hawks, Claudia E. Perez, Sheryl Coutermarsh-Ott, Nisha K. Duggal

## Abstract

Zika virus (ZIKV) can infect developing fetuses *in utero* and cause severe congenital defects. This *in utero* transmission can occurs following ZIKV infection during pregnancy via sexual transmission or mosquito bite. Infected men may shed ZIKV RNA in semen for over six months post symptom onset, indicating that ZIKV may persistently infect the male reproductive tract (MRT). However, the site of persistent infection in the MRT and whether ZIKV can recrudesce in the MRT is unknown. We hypothesized that if ZIKV establishes a persistent infection in the MRT, then immunosuppressant treatment should stimulate ZIKV replication. We tested this hypothesis in a wild-type mouse model of ZIKV sexual transmission. Male mice were infected with ZIKV and immunosuppressed when they no longer shed infectious virus in their ejaculates. After immunosuppression, ejaculates and MRT tissues were monitored for infectious virus and ZIKV RNA. Our results show that ZIKV recrudescence did not occur following immunosuppression, as we did not detect significant levels of infectious virus in ejaculates or MRT tissues following immunosuppression. We did detect ZIKV RNA in the epididymides of mice treated with the immunosuppressant cyclophosphamide. Further analysis revealed that this ZIKV RNA was contained within the lumen of the epididymis. Our findings suggest that ZIKV persistently infects the epididymis within the male reproductive tract. This study provides insight into the mechanisms behind ZIKV sexual transmission, which may inform public health decisions regarding ZIKV risks.

**Importance:** Zika virus (ZIKV) is an emerging mosquito-transmitted virus that typically causes mild and self-limiting febrile illness in humans; however, during the recent epidemic of ZIKV in the Americas, severe birth defects, such as microcephaly and club foot, were reported in infants born to ZIKV infected mothers. Additionally, sexual transmission has been identified as a secondary method of ZIKV transmission. Since ZIKV can be isolated from semen of infected men long after initial infection, it is imperative to understand the mechanism(s) of ZIKV infection of the male reproductive tract to prevent sexual transmission and ZIKV-associated birth defects. The significance of our research is in identifying a site of persistent ZIKV infection in the male reproductive tract and in assessing the likelihood that a persistently infected individual will begin shedding infectious virus in semen again. This information will enhance our understanding of ZIKV sexual transmission and inform health decisions regarding ZIKV risks.

## INTRODUCTION

Zika virus (ZIKV; *Flaviviridae* family, flavivirus genus) can cause severe birth defects, such as microcephaly and club foot, in infants born to mothers infected with ZIKV during pregnancy. These birth defects are collectively termed congenital Zika syndrome and occur in approximately 5-15% of ZIKV-infected pregnancies [1-4]. ZIKV is typically transmitted by the bite of an infected *Aedes* spp. mosquito (*Ae. aegypti or Ae. albopictus*), but sexual transmission of ZIKV was reported during the most recent epidemic [5-9]. Mathematical modelling estimates that sexual transmission accounted for 3-23% of ZIKV transmission in areas with ZIKV-infected mosquitoes [10-12]. Furthermore, *in vivo* in mice studies indicate that maternal ZIKV infection via sexual transmission results in higher viral titers in fetal tissue compared to maternal ZIKV infection via subcutaneous injection, a route of infection that resembles a mosquito bite [13]. Therefore, it is critical that we understand the mechanisms behind ZIKV sexual transmission to reduce ZIKV transmission potential and ultimately prevent serious sequelae, such as ZIKV congenital syndrome.

ZIKV-infected men may shed infectious ZIKV and ZIKV RNA for weeks or even months post infection, potentially increasing the amount of time that they are infectious compared to mosquito transmission [14-18]. Infectious ZIKV has been isolated from the semen of infected men for up to 38 days post onset of symptoms [17]. Additionally, ZIKV RNA has been isolated from semen for over 6 months post symptom onset [18]. Semen is derived from the major internal components of the MRT, which function to produce sperm (testes), mature and store sperm (epididymis) and contribute nutrients, fluids, and other non-cellular components of semen (seminal vesicles and prostate). Acute ZIKV infection of these tissues has been observed in human explant or cell culture models (testes and prostate), mouse models (testes, epididymides, and seminal vesicles), and non-human primate models (testes and prostate) [19-25]. However, persistent ZIKV infection of any of these tissues has yet to be verified.

Persistent infections with flaviviruses are not uncommon. In fact, several of the encephalitic flaviviruses, such as West Nile virus, Japanese encephalitis virus, and tick-borne encephalitis virus, can persistently infect humans, with infectious virus and viral RNA being isolated months to years after the initial symptomatic infection [26-30]. Persistent infections of flaviviruses can be evaluated using *in vivo* models by treating subjects with an immunosuppressant following convalescence and monitoring for viral replication [31-33]. In a mouse model of West Nile virus infection, viral replication was observed in the central nervous system post treatment with the immunosuppressant cyclophosphamide [31].

Our goal in this study was to determine whether ZIKV establishes a persistent infection in the MRT by testing whether immunosuppression could trigger recrudescence of seminal shedding of infectious ZIKV. We used a mouse model of ZIKV sexual transmission that replicates the kinetics of ZIKV shedding in human semen [19, 20]. Following acute infection, male mice were treated with one of a panel of immunosuppressants chosen for their varying mechanisms of action, and viral replication in the MRT was assessed. We were unable to detect infectious virus or an increase in viral RNA in ejaculates following immunosuppression. Low levels of infectious virus were detected in the testes and epididymides in some mice treated with certain immunosuppressants. Lastly, an increase in viral RNA was detected in the epididymides of mice treated with cyclophosphamide. These results suggest that ZIKV infection does establish a persistent infection within the MRT, specifically in the epididymis; however, we were unable to determine whether recrudescence in the MRT can lead to additional shedding of infectious virus in ejaculates.

## METHODS

### Virus strains and cells

Zika virus strain DakAr41524 was used for this study. This strain was isolated in Senegal in 1984 and has since been passaged seven times (AP-61 (*Aedes pseudoscutellaris*) cells p1, C6/36 (*Aedes albopictus*) cells p2, Vero cells p3-7). This strain has been used in studies investigating sexual transmission of ZIKV and can infect mouse testes, epididymides, and seminal vesicles *in vivo*. Additionally, mice infected with this strain shed ZIKV in ejaculates [20].

Vero cells (for plaque assay) were cultured in Dulbecco’s modified Eagle’s medium (DMEM) with 5% fetal bovine serum (FBS), 100units/mL penicillin (Gibco), and 100μg/mL streptomycin (Gibco).

### Mouse inoculations and immunosuppression

Twelve-week-old male C57BL/6J were obtained from the Jackson Laboratory. Mice were allowed to acclimate in an ABSL-2 facility for one day before initial immunosuppression. Mice were rendered susceptible to ZIKV infection via intraperitoneal (i.p.) injection of 2mg of α-IFNAR1 antibody (Mar1-5A3; Leinco Technologies) [20, 34-36]. The following day mice were infected with either 10^3^ (cyclophosphamide) or 10^4^ (remaining drug treatments) PFU of ZIKV via subcutaneous (s.c.) injection in a rear footpad. On days 1 and 4 post infection, mice were given additional doses of 0.5mg of α-IFNAR1 antibody via i.p. injection [20, 35]. Mice were weighed daily to monitor clinical signs of infection. Any mouse whose weight dropped below 85% of the starting weight was humanely euthanized. Blood was collected from the submandibular vein into serum collection tubes on days 3, 5, and 7 post infection. Serum was separated via centrifugation at 10,000 x g for 5 minutes and was stored at -80°C. To collect ejaculate samples, male mice were paired with 1 to 2 CD-1 female mice (Charles River) each night beginning at day 5 post infection. Female mice were checked each morning for evidence of copulation plug. Females who successfully mated were humanely euthanized, and their uteri were dissected out and flushed with BA-1 diluent (1X M199 Hank’s Salts (Sigma), 0.005M Tris-HCL pH7.5 (Gibco), 1% Bovine Serum Albumin (v/v; Probumin; Millipore), 2mM L-glutamine (Gibco), 0.35g/L Sodium Bicarbonate (Gibco), 100 units/mL Penicillin (Gibco), 100ug/mL streptomycin (Gibco), and 1ug/mL Amphotericin B (Hyclone)) to collect the ejaculate.

After male mice cleared the initial infection, as evidenced by weight gain and lack of infectious ZIKV in serum and ejaculate samples (via plaque assay), mice were immunosuppressed via cyclophosphamide, dexamethasone, ketoconazole/cyclosporine, methylprednisolone acetate, or α-IFNAR1 antibody. Cyclophosphamide (Sigma), dissolved in PBS, was administered at 5mg/mouse via i.p. injection on days 31 and 36 post infection. Water-soluble dexamethasone (Sigma), dissolved in PBS, was administered at 1mg/kg via oral gavage daily from dpi 32-42. Ketoconazale (Sigma), dissolved in peanut oil (Sigma), was administed at 10 mg/kg via oral gavage daily from dpi 32-42. Cyclosporine (Sigma), dissolved in DMSO (ATCC) and diluted in PBS, was administered at 30 mg/kg via i.p. injection daily from dpi 32-42. Methylprednisolone acetate (Zoetis) was administered at 600 mg/kg via s.c. injection on the back on day 32 post infection. α-IFNAR1 antibody, diluted in PBS, was administered at 2 mg per mouse via i.p injection on day 32 post infection and 0.5 mg per mouse via i.p. injection on days 34 and 37 post infection. Mice infected with either 10^3^ or 10^4^ PFUs that received PBS via i.p. injection on days 32, 34, and 37 post infection served as controls.

Male mice were humanely euthanized ten days after immunosuppression. Blood was collected via submandibular vein or intracardiac bleed. Serum was separated and stored as described above. Testes, epididymides, and seminal vesicles were dissected out of each mouse. One set of reproductive tissues from each mouse was preserved in neutral buffered formalin for later ISH and H&E analysis, while remaining tissues were frozen at -80°C for later virus quantification.

### Quantification of infectious virus

Infectious virus was quantified via Vero cell plaque assay. Briefly, tissues from mice were weighed, and an equal volume of BA-1 diluent was added to each sample. One 5mm stainless steel bead was added to each sample, and tissues were homogenized using a TissueLyserLT (Qiagen) set to 5 oscillations/s for 2 minutes. Tissue samples were clarified via centrifugation at 19,000g for 3 minutes.

Serum, ejaculate, and clarified tissue samples were serially diluted in BA-1 diluent. These dilutions were plated on confluent Vero cells and incubated at 37°C, 5% CO_2_ for 1 hour, with gentle rocking every 15 minutes. Following incubation, Vero cells were overlaid with Miller’s Ye-Lah agarose overlay (2X Ye-Lah media (0.132% yeast extract (w/v), 0.66% lactalbumin hydrolysate (w/v), 10X Earle’s Balanced Salt Solution, 2% Fetal Bovine Serum (v/v), Amphotericin B, Gentamycin), 1.6% agarose (w/v), and 0.225% sodium bicarbonate (v/v)). A second overlay containing neutral red (1:300) was added four days later. Plaques were counted the following day. The limit of detection is 2 log_10_ PFU/mL serum, 0.4 log_10_ PFU/ejaculate, and 0.4 log_10_ PFU/organ.

### Quantification of viral RNA

Viral RNA was extracted from ejaculates and homogenized tissue samples using the QIAamp viral RNA mini kit (Qiagen). Briefly, 70µL sample was diluted 1:2 in 10µM dithiothreitol (DTT; Pierce) to denature seminal proteins. Samples were lysed in 560µL buffer AVL with linear acrylamide added (1µg per sample). The extraction was continued following the manufacturer’s protocol. Viral RNA was eluted in 60µL nuclease-free water (Qiagen) and stored at -80C.

Viral RNA was quantified via a one-step qRT-PCR using the iTaq universal probes one-step kit (Bio-Rad) per the manufacturer’s instructions for a 20µL reaction, with the exception that the quantity of reverse transcriptase per reaction was halved. Five microliters of RNA were used per reaction. ZIKV specific primers and probes were synthesized by IDT using 6-Fam as the reporter dye and Zen/Iowa Black as the quencher. Primer and probe sequences and cycling conditions are as previously described [37]. The amplification product is an approximately 75bp region of the envelope protein. Viral RNA concentration was determined via an absolute standard curve of *in vitro* transcribed RNA standards from a plasmid containing a segment of the ZIKV envelope gene [20, 37]. The limit of detection for this assay was 1 RNA copy per reaction, 2 log_10_ RNA copies per ejaculate, or 1.5 log_10_ RNA copies per organ.

### Visualization of viral proteins within tissues via immunohistochemistry (IHC)

Testes, epididymides, and seminal vesicles were collected from male mice upon euthanasia and fixed and stored in 10% buffered formalin. Tissues were paraffin-embedded, and 5μM slices were attached to charged, glass slides. Tissues were deparaffinized by submersion of slides in Xylenes (Fisher Scientific) followed by submersion in decreasing concentrations of ethanol (Fisher Scientific). Antigen retrieval was performed by submersion in 10mM sodium citrate buffer at 91 to 95°C for 30 minutes. After cooling, tissues were stained using the Peroxidase IHC Detection Kit (Pierce), per the manufacturer’s protocol. Primary antibody was anti-ZIKV NS2B (GeneTex GTX133308) and secondary antibody was goat anti-rabbit conjugated to horseradish peroxidase (Invitrogen). Both antibodies were diluted 1:500 in universal blocking buffer (Pierce) before use. Positive staining (brown) was detected using DAB substrate (Pierce). Nuclei were counterstained (positive) using Harris-modified hematoxylin (Pierce) for two minutes. Tissues from uninfected mice served as negative controls, while tissues from mice who succumbed to ZIKV were used as positive controls.

### Visualization of viral RNA within tissues via In Situ Hybridization (ISH)

Tissues were collected and processed as for IHC. ISH was performed using the view RNA ISH Tissue Assay (Invitrogen) per the manufacturer’s instructions. Tissues underwent pretreatment and protease treatment for 10 minutes each [38]. A proprietary but publicly available probe set specific for positive sense ZIKV (Asian lineage) RNA was used (Thermofisher). Positive staining (red) was visualized via alkaline phosphatase labeled probes. Nuclei (blue) were counterstained via Gill’s hematoxylin (American Master Tech Scientific) for 3 minutes. Tissues from uninfected and infected mice euthanized during acute infection served as negative and positive controls, respectively.

### Histological analysis

Tissues were collected and processed as for IHC. Slides were stained for Hematoxylin and Eosin (H&E) following normal procedures. Slides from infected mice euthanized before immunosuppression (dpi 31) and from uninfected mice treated with cyclophosphamide or PBS served as controls. Slides were analyzed by Sheryl Coutermarsh-Ott, DVM, PhD, Diplomate of the American College of Veterinary Pathologists (DACVP). Testes were assessed for degradation of tubule architecture, inflammation of interstitial spaces, and Leydig cell loss. Epididymides were assessed for epithelial damage and interstitial inflammation. Each of these factors were scored from 0 (no pathology) to 3 (severe pathology), and the scores from each of these subcategories were added together to achieve a total organ score.

### Statistics

Statistical analyses were performed using GraphPad Prism (v8.4.1). Weight data were analyzed using repeated measures analysis of variance (ANOVA) with multiple comparisons t-tests using Tukey correction. Infectious virus and viral RNA concentrations in ejaculates were assessed via multiple comparisons t-tests using the Holm-Sidak correction. Correlations were assessed via Spearman correlation coefficient (ZIKV RNAc in epididymis vs. peak ZIKV RNAc in the ejaculates or number of matings), Mann-Whitney rank sum test (ISH results vs. ZIKV RNAc in epididymis, number of matings, viremia at 3 dpi, or viremia at 5 dpi).

### Ethics statement

All animal experiments were approved by the Institutional Animal Care and Use Committee at Virginia Polytechnic Institute and State University (IACUC protocol 18-085) and followed the recommendations in the *Guide for the Care and Use of Laboratory Animals*, 8^th^ edition (Institute for Laboratory Animal Research, National Research Council, National Academy of Sciences, 2011).

## RESULTS

### Immunosuppression following ZIKV infection does not lead to systemic recrudescence

To assess whether ZIKV can recrudescence in the MRT, C57BL/6J male mice pre-treated with an IFNAR blocking antibody were infected subcutaneously with ZIKV strain Dakar41524. Serum and ejaculates were monitored regularly for presence of infectious virus. When infectious virus was no longer shed in ejaculates (∼30 dpi), male mice were treated with one of the following immunosuppressants chosen for their differing mechanisms of action: cyclophosphamide, IFNAR blocking antibody, methylprednisolone acetate, dexamethasone, or ketoconazole/cyclosporine. PBS treated mice served as a control. Post-immunosuppression, serum and ejaculates were collected regularly to assess for infectious virus. Mice were euthanized ten days post immunosuppression, and testes, epididymides, and seminal vesicles were collected and assessed for the presence of infectious virus (Figure 1a).

**Figure 1:**
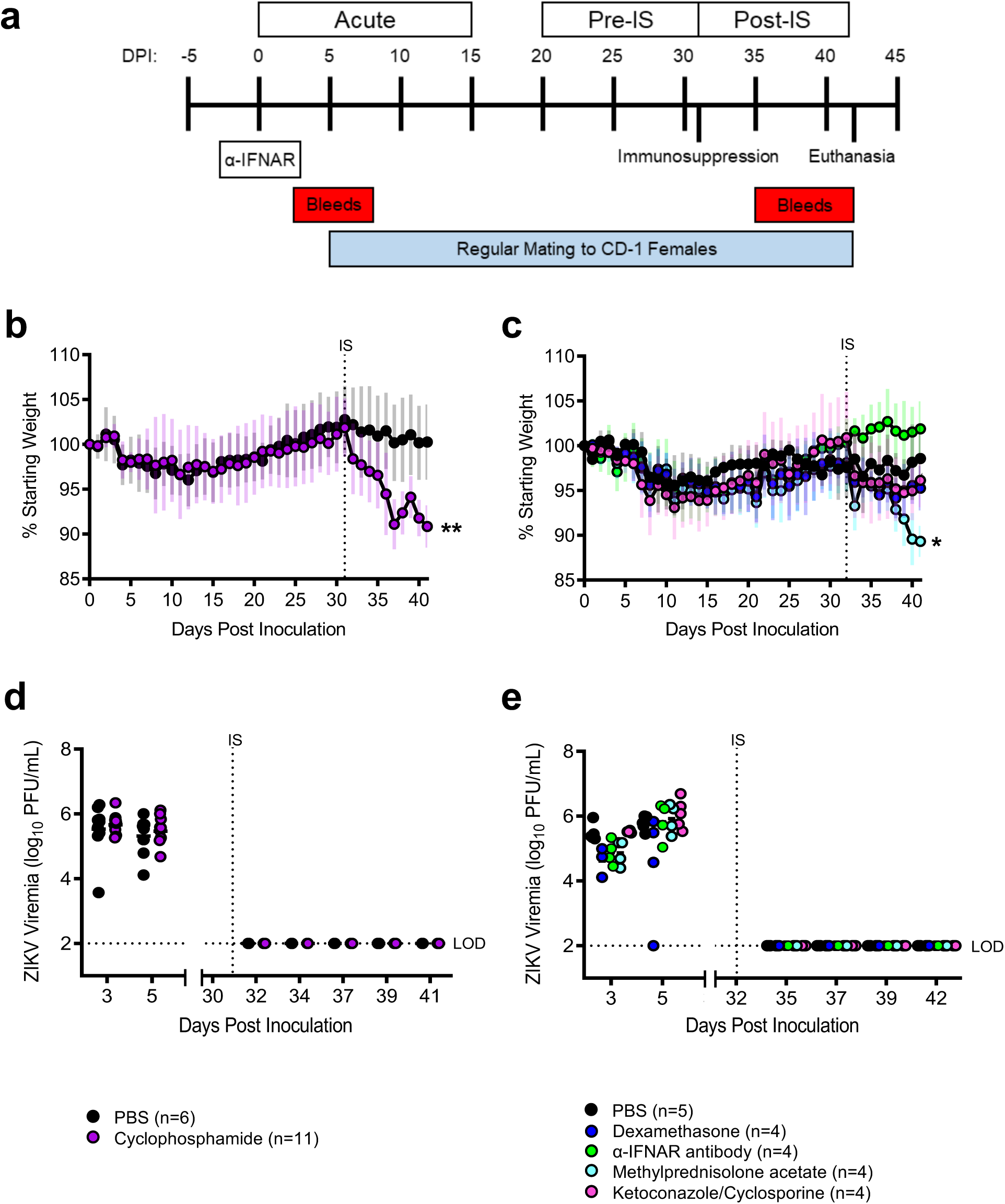
Treatment with immunosuppressants affects morbidity but not mortality or viremia in ZIKV-infected male mice. Study design (**a**). Mouse weights were recorded daily (mean ± standard deviation) PBS: n=6; Cyclophosphamide (**b)**. Mouse weights PBS: n=5; Dexamethasone: n=4; α-IFNAR antibody: n=4; Methylprednisolone acetate: n=4; Ketoconazole/Cyclosporine: n=4 (**c**). Infectious ZIKV in serum was quantified via plaque assay (**d**,**e**). Data points represent individual mice, horizontal lines represent group mean, and error bars represent standard deviation. Data were analyzed via repeated measures ANOVA. * p<0.05; ** p<0.01; *** p<0.005; **** p<0.001. Abbreviations: DPI, days post inoculation; IS, immunosuppression; LOD, limit of detection.

Mortality, morbidity, and viremia were monitored throughout the course of the study. During acute infection, there was a 20% mortality rate in mice infected with 10^3^ PFUs of ZIKV and a 24% mortality rate in mice infected with 10^4^ PFUs of ZIKV, with mortalities occurring between dpi 8 and 10. No mortalities occurred post-immunosuppression, regardless of the immunosuppressant used. On average, mice lost approximately 5% of their starting weight during acute infection and gained that weight back during the pre-immunosuppression phase (Figures 1B and 1C). Post-immunosuppression, there was significant weight loss in mice treated with cyclophosphamide (10% weight loss; p = 0.003) and methylprednisolone acetate (9% weight loss; p=0.03) compared to their respective PBS treated controls (2% weight loss); however, it is likely that this weight loss was due to the immunosuppressant agents themselves and not due to ZIKV recrudescence, no infectious virus was detected in serum post-immunosuppression, regardless of immunosuppressant treatment (Figures 1d and 1e). In contrast, infectious virus was present in serum at concentrations ranging from 3.6 to 6.7 log_10_ PFUs/mL during acute infection. Taken together, these results confirm that mice were infected with ZIKV experienced acute disease but did not develop systemic ZIKV recrudescence following immunosuppression.

### Immunosuppression does not increase concentration ZIKV in ejaculates of ZIKV infected male mice

To assess whether ZIKV recrudescence occurred in the MRT following immunosuppression, infectious virus was quantified in ejaculates via plaque assay (Figures 2a and 2b). During acute infection, male mice shed up to 6 log_10_ PFU in ejaculates, with infectious virus cleared by twenty days post inoculation. No infectious virus was detected in ejaculates post immunosuppression regardless of immunosuppressant treatment. Next, ZIKV RNA levels were quantified in ejaculates via qRT-PCR (Figures 2c and 2d). ZIKV RNA was detected in ejaculates throughout the course of the entire study with the highest concentrations (5 to 6 log_10_) detected in samples collected during acute infection. To determine whether ZIKV RNA concentrations increased in ejaculates post-immunosuppression, samples were grouped based on collection time: 10 days pre-immunosuppression and 10 days post-immunosuppression (Figures 2G and 2H). ZIKV RNA levels in ejaculates from mice immunosuppressed with the α-IFNAR antibody were significantly higher post-immunosuppression than those from pre-immunosuppression samples (p=0.03). No other immunosuppressant treatments increased ZIKV RNA in ejaculates. There were no significant changes in ZIKV RNA levels in ejaculates pre- and post-immunosuppression in PBS treated mice. Taken together, these data indicate that immunosuppression did not significantly increase infectious ZIKV in mouse ejaculates.

**Figure 2:**
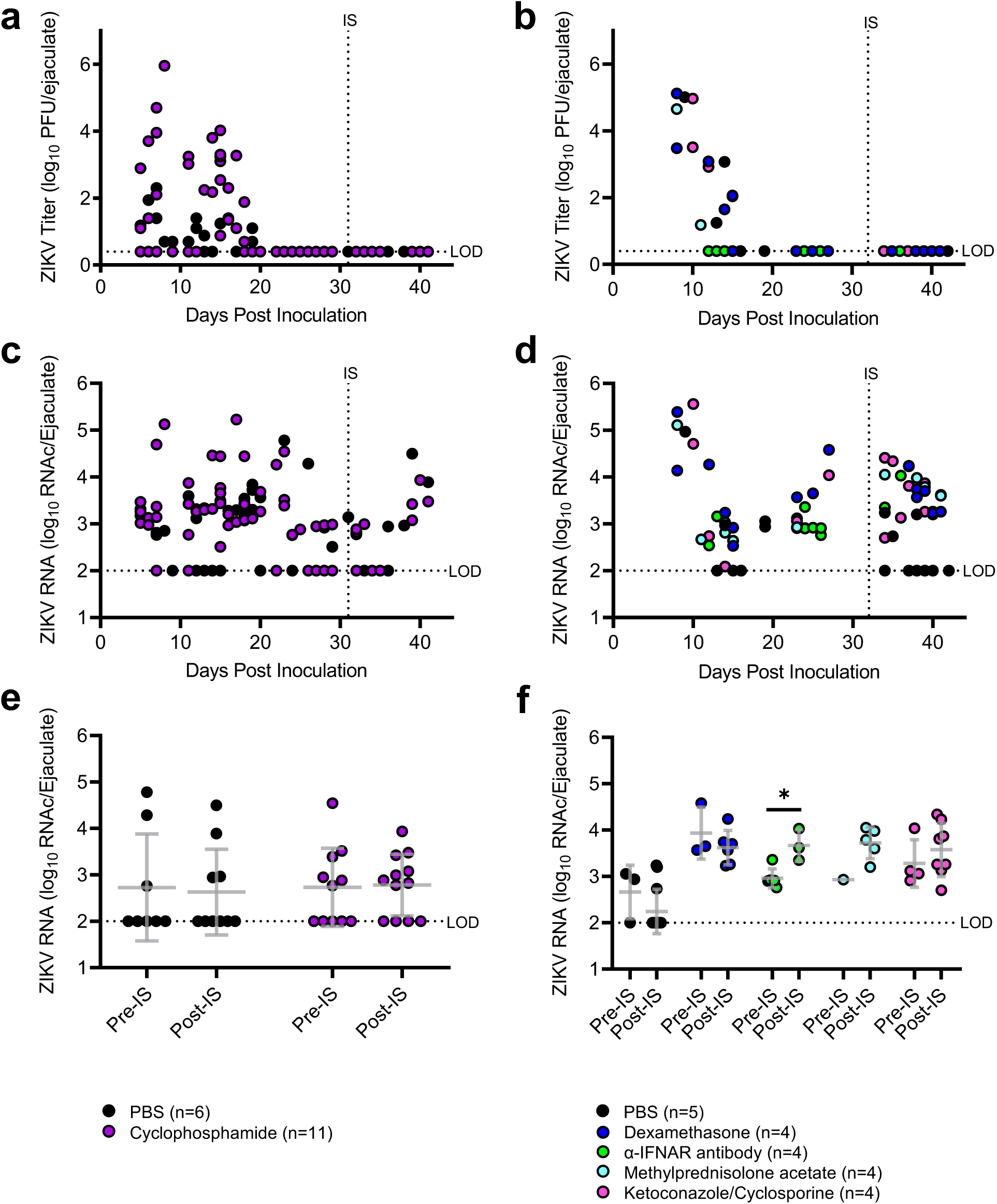
ZIKV RNA but not infectious ZIKV are present in ejaculates from ZIKV-infected male mice treated with immunosuppressants. Infectious ZIKV in ejaculate samples was quantified via plaque assay **(a**,**b)**, and ZIKV RNA was quantified via qRT-PCR **(c**,**d)**. For further analysis of qRT-PCR results, ejaculate samples were grouped based on the stage of infection in which they were collected **(e**,**f)**: pre-immunosuppression (days 22-27) and post-immunosuppression (days 33-42). Each data point represents an individual sample, with lines and error bars representing the mean and standard deviation, respectively. Data were analyzed via two-way ANOVA followed by multiple comparisons t-tests using the Holm-Sidak correction. * p<0.05; ** p<0.01; *** p<0.005; **** p<0.001. Abbreviations: RNAc, RNA copies; LOD, limit of detection; IS, immunosuppression; ns, not significant.

### Cyclophosphamide treatment significantly increases ZIKV RNA but not infectious virus in epididymides

Male reproductive tract tissues (testes, epididymides, and seminal vesicles) were collected from mice upon euthanasia, and infectious virus was quantified via plaque assay. Infectious virus was detected in reproductive tract tissues of mice who succumbed to acute ZIKV infection, confirming that ZIKV does infect MRT tissues during acute infection. Infectious virus was not detected in tissues from any of the mice treated with cyclophosphamide or PBS (Figures 3a and 3c). Infectious virus was detected at or near the limit of detection in the testes of one mouse treated with dexamethasone and the epididymides of two mice treated with dexamethasone and two mice treated with methylprednisolone acetate (Figures 3b and 3d). We confirmed these results via immunohistochemistry using an antibody against ZIKV NS2B protein, and we were able to detect NS2B in 3 out of the 7 samples that were positive via plaque assay.

**Figure 3:**
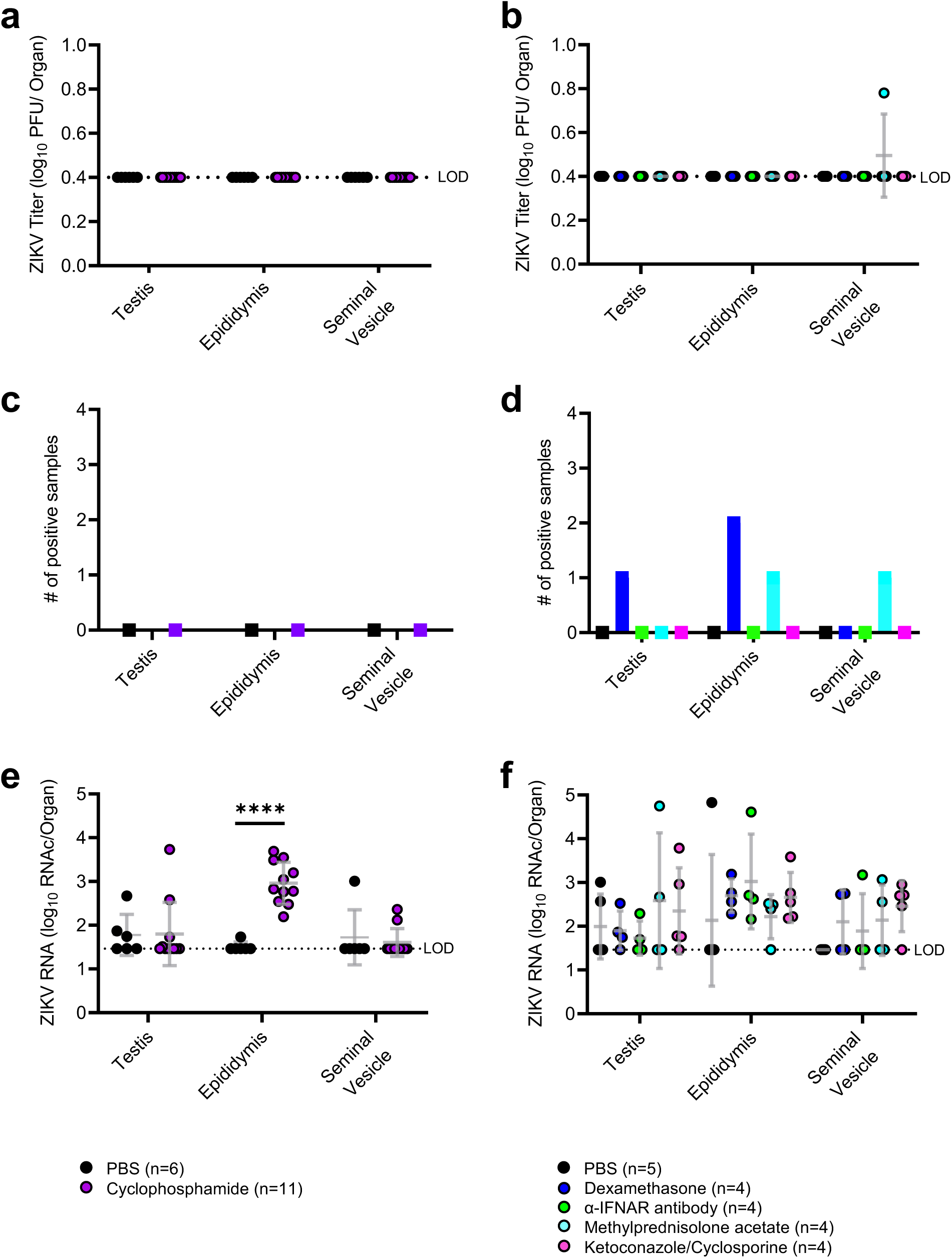
Infectious ZIKV and ZIKV RNA are present in male reproductive tract tissues of ZIKV-infected mice following treatment with select immunosuppressants. Male reproductive tract tissue samples were collected from mice upon euthanasia (day 41-42). Tissues were homogenized, and infectious ZIKV was quantified via plaque assay **(a**,**b)**. The number of samples that yielded positive plaque assay results were counted **(c**,**d)**. ZIKV RNA copies were quantified via qRT-PCR **(e**,**f)**. Each data point represents an individual sample, with lines and error bars representing the mean and standard deviation, respectively. Data were analyzed via two-way ANOVA followed by multiple comparisons t-tests using the Holm-Sidak correction. * p<0.05; ** p<0.01; *** p<0.005; **** p<0.001. Abbreviations: RNAc, RNA copies; LOD, limit of detection; ns, not significant.

Levels of ZIKV RNA in MRT tissues were assessed via qRT-PCR (Figures 3e and 3f). ZIKV RNA was significantly higher in epididymides from mice treated with cyclophosphamide than from those treated with PBS (p <0.001). There were no significant changes in ZIKV RNA levels in tissues of mice treated with any of the other immunosuppressants compared to PBS controls. To confirm the presence of ZIKV RNA in the epididymides of cyclophosphamide treated mice, we performed *in situ* hybridization (ISH) against ZIKV RNA in the epididymides of cyclophosphamide and PBS treated mice (Figure 4). We detected ZIKV RNA via ISH in the epididymal lumen of 5 of the cyclophosphamide treated mice (n=10) but in none of the epididymides from the PBS treated mice (n=3). To ascertain whether ZIKV was actively replicating in these epididymides, we performed IHC against NS2B protein, but we were unable to detect ZIKV NS2B protein in epididymides with positive ISH results. Additionally, we performed correlation analyses to determine whether the positive ISH results were related to ZIKV titers throughout infection or other variables. There were no significant correlations between ZIKV RNAc in the epididymis and number of matings (p=0.33), ISH results (p=0.23), or ZIKV RNAc in the ejaculates (p=0.15), as well as no significant correlation between ISH results and number of matings (p=0.94), viremia at 3 dpi (p=0.27), or viremia at 5 dpi (p=0.072).

**Figure 4:**
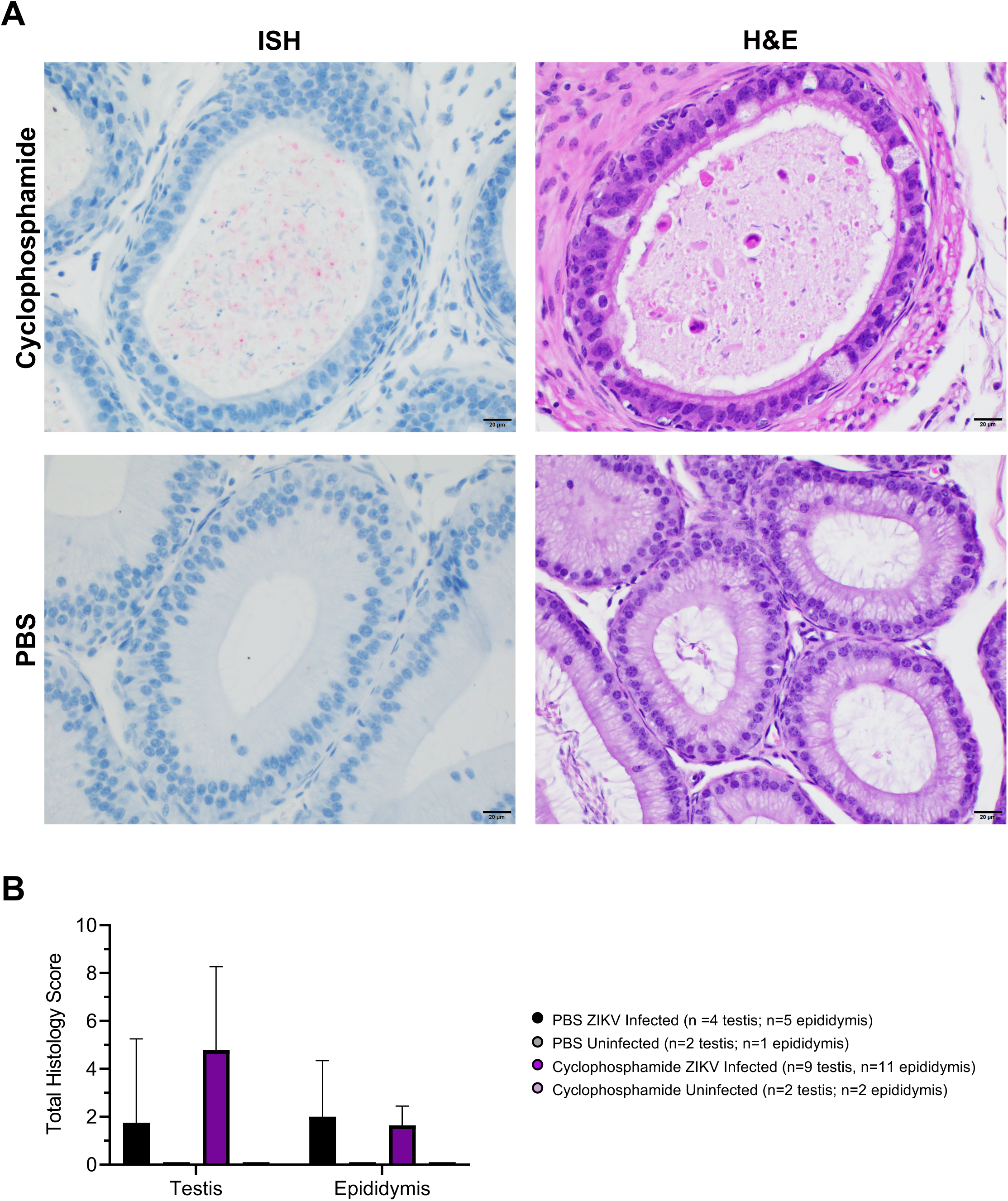
ZIKV RNA is present within the lumen of epididymides of cyclophosphamide treated mice. Testes and epididymides from cyclophosphamide and PBS treated mice were assessed for ZIKV RNA via ISH. ZIKV RNA is stained red and nuclei are stained blue. The same tissues were also stained via H&E to assess histology Five out of 10 epididymides from cyclophosphamide mice and 0 out of 3 from PBS mice had positive ISH staining **(a)**. H&E slides were scored for total testicular or epididymal pathologies. Data are displayed as mean with error bars representing standard deviation **(b)**. Statistical analyses were assessed using the Mann-Whitney ranked sum test. * p<0.05; ** p<0.01; *** p<0.005; **** p<0.001.

Sections of testis and epididymis were evaluated histologically for evidence of tissue damage and inflammation. In general, all ZIKV mice exhibited some degree of tissue pathology. In the testis, these changes ranged from moderate increases in interstitial lymphocytes and plasma cells to massive loss of seminiferous tubules with collapse of normal architecture. In the epididymis, these changes were less severe with mild infiltration of inflammatory cells and mild to moderate degeneration and loss of epithelial cells. These changes were graded semi-quantitatively to produce a total histologic score. There were no significant differences in histologic scores of testes (p=0.082) or epididymides (p=0.86) between cyclophosphamide and PBS treated ZIKV infected mice (Figure 4). No pathology was observed in testes or epididymides of uninfected mice treated with PBS or cyclophosphamide, indicating that cyclophosphamide treatment alone does not cause MRT pathologies. Severe testicular damage and mild to moderate epididymal damage were observed in tissues from mice euthanized immediately before immunosuppressant treatment, indicating that MRT pathology may have manifested before immunosuppressant treatment. Taken together, these results indicate that cyclophosphamide treatment increased ZIKV RNA in the epididymides, specifically in non-sperm cells within the lumen, and that cyclophosphamide treatment did not impact MRT tissue pathology.

## DISCUSSION

ZIKV infection in pregnant women can lead to severe congenital defects in the developing fetus. *in utero* ZIKV transmission can occur if the mother is infected via sexual or mosquito bite during pregnancy [39]. Since ZIKV RNA has been detected in human semen for six months post symptom onset [14, 15, 17, 18], we investigated whether ZIKV persists in the MRT and could recrudescence upon immunosuppressant treatment in a mouse model. We observed that immunosuppression did not stimulate systemic ZIKV recrudescence or resumption of shedding of infectious ZIKV in ejaculates. However, ZIKV RNA levels as detected by qRT-PCR were significantly increased in the epididymides of mice treated with the immunosuppressant cyclophosphamide compared to PBS-treated controls. Additionally, ZIKV RNA was visualized via ISH in in the extracellular, luminal contents of the epididymis of mice treated with cyclophosphamide. Rarely, we also identified them within degenerate cells within epididymis of these mice as well. Collectively, these results indicate that ZIKV infection may persist within the epididymis.

In humans, infectious ZIKV and ZIKV RNA have been isolated from ejaculates up to 38 days and 370 days post symptom onset, respectively [17, 18, 40]. While this long period of ZIKV RNA shedding is likely a result of persistent infection in the MRT, the source of the ZIKV RNA in the MRT and the potential for further transmission are unknown. We hypothesized that immunosuppression after the acute phase of ZIKV infection would stimulate ZIKV replication in reservoirs within the MRT, if any exist. This method allowed us to identify the epididymis as a potential site for persistent ZIKV infection. In mouse models of ZIKV infection, the testes atrophy and sustain severe pathologies including loss of seminiferous tubule structures and interstitial cell populations [41-43], though these testicular pathologies have not been observed in human cases of ZIKV or in *ex vivo* ZIKV infections of human testicular explants [21]. We did observe testicular atrophy and pathology in mice in our study, and it is possible that ZIKV susceptible cells in these testes were eliminated due to these pathologies, preventing establishment of a persistent infection in the testes. To better assess whether ZIKV establishes a persistent infection in the testes, a model organism that does not experience severe testicular pathologies following ZIKV infection, such as non-human primates, might need to be used [25, 44]. However, our results indicate that immunosuppression is unlikely to result in the recurrence of infectious ZIKV in ejaculates. Following immunosuppression, we were unable to detect infectious ZIKV in ejaculates; however, we only assessed ejaculates for ten days post-immunosuppression. In a study of persistence of West Nile virus in a mouse model, recrudescence was detected in tissues fifteen days post cyclophosphamide treatment [31]. We were unable to investigate ZIKV recrudescence in ejaculates fifteen days post treatment because the collection of ejaculates became difficult. Given that many of the mice lost weight following immunosuppression, especially those treated with cyclophosphamide or methylprednisolone acetate, we speculated that immunosuppression caused the mice to feel unwell, which reduced mating.

The infection of the epididymis during acute ZIKV infection has been well characterized in a mouse model [38]. Infection begins in the head of epididymis and quickly spreads to the tail of the epididymis. Epididymal epithelial cells and luminal leukocytes were found to be the targets of ZIKV infection during acute infection. In our study, we did find ZIKV RNA in the epididymis, but it was primarily extracellular and only within luminal contents. While the epididymal epithelium was damaged, there was no evidence of epididymal epithelium infection post immunosuppression. Given that the epididymis functions, in part, as a storage for mature sperm before ejaculation, it is possible that the infected luminal cells we observed were not a result of persistent ZIKV infection but rather residual infected cells that had yet to clear the MRT. In humans, duration of ZIKV shedding in semen is inversely correlated with the frequency of ejaculation [18]. In our study, we found that there was no correlation between mating frequency and epididymal ZIKV RNA levels by qRT-PCR or ISH staining. Additionally, there were no correlations between ZIKV ISH staining and viremia titers, peak ejaculate RNA copies, or epididymal RNA copies. These results indicate that ZIKV RNA in the epididymis is more likely a result of persistent ZIKV infection as opposed to residual infected cells in the epididymis. We attempted to validate this conclusion by performing IHC on these tissues for the viral protein NS2B, which is only present in cells with actively replicating ZIKV; however, we were unable to detect any NS2B in our samples.

Interestingly, we did identify ZIKV RNA rarely within cells within the lumen of epididymis of immunosuppressed mice. These cells were often degenerate and thus unable to be definitively identified solely on morphology. It is suspected, however, that these cells are likely macrophages or degenerate epithelial cells. Macrophages are present throughout the MRT in both the interstitial spaces of the testes and epididymis; however, macrophages rarely cross the blood testes barrier or the blood epididymis barrier (which serve to maintain immune-privileged sites for sperm development and maturation) in healthy individuals [45-47]. ZIKV infection may disrupt these barriers in the MRT, allowing for macrophages to enter the seminiferous tubules of either the testes or the epididymis [48, 49]; however, it is unknown whether macrophages are infected before or after entrance into these tubules. Degenerate epithelial cells are derived from epididymal epithelial cells that are sloughed into the lumen upon epithelial damage. Since ZIKV does infect the epithelium of the epididymis and causes epithelial damage [38, 50], infected degenerate epithelial cells within the epididymal lumen were likely infected before being sloughed off into the lumen. More work will need to be done to determine how ZIKV reaches the epididymis and establishes persistence. Additionally, we did not detect ZIKV RNA within sperm cells in the epididymis, indicating that a male with a persistent ZIKV infection may be able to safely conceive a child using *in vitro* fertilization or similar reproductive technologies where sperm cells are separated from the remainder of the semen.

The mechanisms of action of the various immunosuppressants used in this study may provide insights into how ZIKV establishes a persistent infection in the MRT and what immune responses are necessary to clear ZIKV from the MRT. The immunosuppressants used in this study were chosen for their differing mechanisms of action, which are as follows: (1) cyclophosphamide induces apoptosis of rapidly dividing cells [31, 51]; (2) α-IFNAR1 antibody inhibits the interferon response by preventing type 1 interferons from binding to their cognate receptors [34]; (3) the combination of ketoconazole and cyclosporine inhibits T cell activation [52-54]; (4) dexamethasone induces apoptosis of peripheral T cells and induction of an anti-inflammatory response [55-57]; and (5) methylprednisolone acetate decreases T cell and monocyte populations and induces an anti-inflammatory response [58]. In our study, we observed an increase in ZIKV RNA only in the epididymides of mice treated with cyclophosphamide. This implies that rapidly dividing cells (such as developing monocytes or macrophages or expanding T or B cell clonal populations) are important in the immune response to ZIKV in the MRT. Additionally, this indicates that peripheral T cells may not be important in the immune response to ZIKV in the MRT. Future studies will delve into the immune response to ZIKV in the MRT, specifically the epididymis.

The long-term impacts of ZIKV on the MRT are still largely unknown. The study presented here provides insights into the role of the epididymis in ZIKV infection and the mechanism of ZIKV persistence in the MRT. Additionally, this study provides the foundation for future studies on immune responses to viral infection in the MRT, particularly within the epididymis. Understanding how ZIKV infects and persists within the MRT may help explain the mechanisms behind ZIKV sexual transmission, allowing for increased knowledge of ZIKV transmission risk and reduced incidence of ZIKV congenital syndrome.

## ACKNOWLEDGEMENTS

Funding for this project was provided by the NIAID (R21A142504). We thank VT Laboratory Animal Research staff for contributions to animal husbandry and training for mouse procedures. We thank the VT ViTALS lab for processing tissues for microscopy and performing H&E staining.

